# Monitoring quinolone resistance due to *Haemophilus influenzae* mutations (2012–17)

**DOI:** 10.1101/632869

**Authors:** Yasuhiro Nagatomo, Tetsuro Shirakura, Kunihiko Fukuchi, Takahiro Takuma, Issei Tokimatsu, Yoshihito Niki

## Abstract

Among drug-resistant bacteria of recent concern, we determined minimum inhibitory concentrations (MICs) of six different quinolone antibacterial agents in *Haemophilus influenzae* and performed molecular genetic analysis in addition to the exploration for β-lactamase-producing and β-lactamase negative ampicillin-resistant *H. influenzae* (BLNAR). A total of 144 clinical *H. influenzae* strains isolated at the Showa University Hospital between 2012 and 2017 were subjected to MIC determination for penicillin/quinolone antibacterial agents using the nitrocefin method and the Clinical and Laboratory Standards Institute broth microdilution method. Moreover, amino acid mutations in the quinolone resistance-determining regions (QRDRs) were analyzed in the isolates showing MIC value of ≥ 0.25 µg/ml of quinolone antibacterial agents. Increasing proportions of BLNAR were noted, with 15% in 2015 to 43.5% in 2016 and 63.6% in 2017. Among quinolone antibacterial agents, all isolates remained susceptible to sitafloxacin (STFX), and STFX showed strong inhibitory potencies against both DNA gyrase and topoisomerase IV. For moxifloxacin (MXF), however, strains with MIC value of 0.5 µg/ml were detected every year since 2013 except 2015. Amino acid mutations were investigated in 17 isolates (11.8%) with MXF MIC value of ≥0.25 µg/ml, and confirmed in 11 isolates (7.6%), of which mutations of GyrA were found in 9 isolates. Future antibacterial drug regimens may need to address the emergence of quinolone-resistant *H. influenzae*.

## Introduction

*Haemophilus influenzae* is the second most common causative bacterium of community-acquired pneumonia, following *Streptococcus pneumoniae*, and has taken the leading position after pneumococcal vaccines became popular (1). β-Lactam antibiotics have previously been used for the treatment of community-acquired pneumonia. In the 2000s, however, β-lactamase producing ampicillin-resistant (BLPAR) bacteria have been increasingly detected in Canada (2), Japan, and some other countries. In addition, β-lactamase negative ampicillin-resistant (BLNAR) bacteria have been increasingly detected in Japan in the recent years (3). Moreover, BLNAR bacteria exhibit reduced susceptibility to cephems, which have high affinity to penicillin binding protein (PBP) 3, thereby raising a concern regarding the possible acquisition of drug resistance.

Meanwhile, fluoroquinolones are clinically effective against these bacteria (4), and in particular, have been frequently prescribed to outpatients with respiratory and otolaryngologic infections. Accordingly, reports on *H. influenzae* resistant to quinolone antibacterial agents have started emerging in the beginning of 2000. In Spain, a report has documented that ciprofloxacin (CIP)-insensitive strains accounted for 0.26% and 0.36% in 2005–2009 and 2010–2013, respectively (5), whereas another report has stated that low-sensitive strains to CIP accounted for 43% in 2007 (6). In Taiwan, the levofloxacin (LVX) resistance rate has been reported to have increased from 2.0% in 2004 to 24.3% in 2010 (7). These data illustrate that the prevalence of quinolone resistance among *H. influenzae* isolates is highly variable depending on when the test is conducted even within the same country or within the same institution. Furthermore, high-level BLNAR strains have been reported to exhibit reduced sensitivity toward fluoroquinolones (8). Thus, it is essential to take into consideration the increasing number of BLNAR isolates and determine the trend of quinolone sensitivity.

Fluoroquinolone resistance is primarily induced through mutations in QRDRs of GyrA gene encoding DNA gyrase or ParC gene encoding topoisomerase IV, which cause three-dimensional structural alterations in the respective enzymes and decrease the affinity between fluoroquinolones and these replication enzymes (9). In the case of *H. influenzae*, drug resistance has been shown to be strongly related to mutations of serine and aspartic acid at positions 84 and 88 of GyrA and of serine at position 84 of ParC (10). However, only a limited number of previous studies have investigated mutations in QRDRs in the same institution over time or have compared MICs among different quinolones. In this study, we tested sensitivity levels to quinolone and penicillin antibacterial agents over 6 years in *H. influenzae* strains that were isolated at the same institution without effects from clinical departments, diseases, or specimens, and analyzed genetic mutations in QRDRs in strains with MIC value of ≥0.25 µg/ml of quinolones. Next, we assessed whether antibacterial efficacies varied among quinolone antibacterial agents or whether the bacteria tended to be resistant.

## Results

### BLNAR and β-lactamase-producing bacteria

We tested the drug susceptibility using 144 *H. influenzae* strains stored in our laboratory. *H. influenzae* strains had been isolated from respiratory (n = 130), otolaryngologic (n = 6), blood (n = 3), pus (n = 3), and other (n= 2) samples. As per the Clinical and Laboratory Standards Institute guidelines, the tests were performed by the broth microdilution method using plates (11). The test bacterial strain was cultured on chocolate agar medium at 35°C in 5% CO_2_ for 22 hours (8). The bacterial strains were seeded in each well at a final inoculum of 5 × 10^5^ CFU/ml. The plates were incubated at 35°C in 5% CO_2_ for 22 hours, and MIC values of the test drugs against clinical *H. influenzae* isolates were measured. Furthermore, the β-lactamase producing ability was determined by the nitrocefin method (12). Based on these results, percentages of BLNAR and β-lactamase-producing bacteria over time are shown in Figure 1. Among all tested strains, BLNAR strains accounted for 38.9% in 2012, 22.2% in 2013, 37.0% in 2014, 15.0% in 2015, 43.5% in 2016, and 63.6% in 2017, and the percentage of BLNAR strains was found to have consecutively increased in 2016 and 2017. In 2017, aminobenzylpenicillin (ABPC)-resistant strains, including β-lactamase-producing ones, accounted for 78.8% of all isolates. β-Lactamase producing amoxicillin/clavulanic acid resistant *H. influenzae* (BLPACR) was not detected.

**Figure 1.**
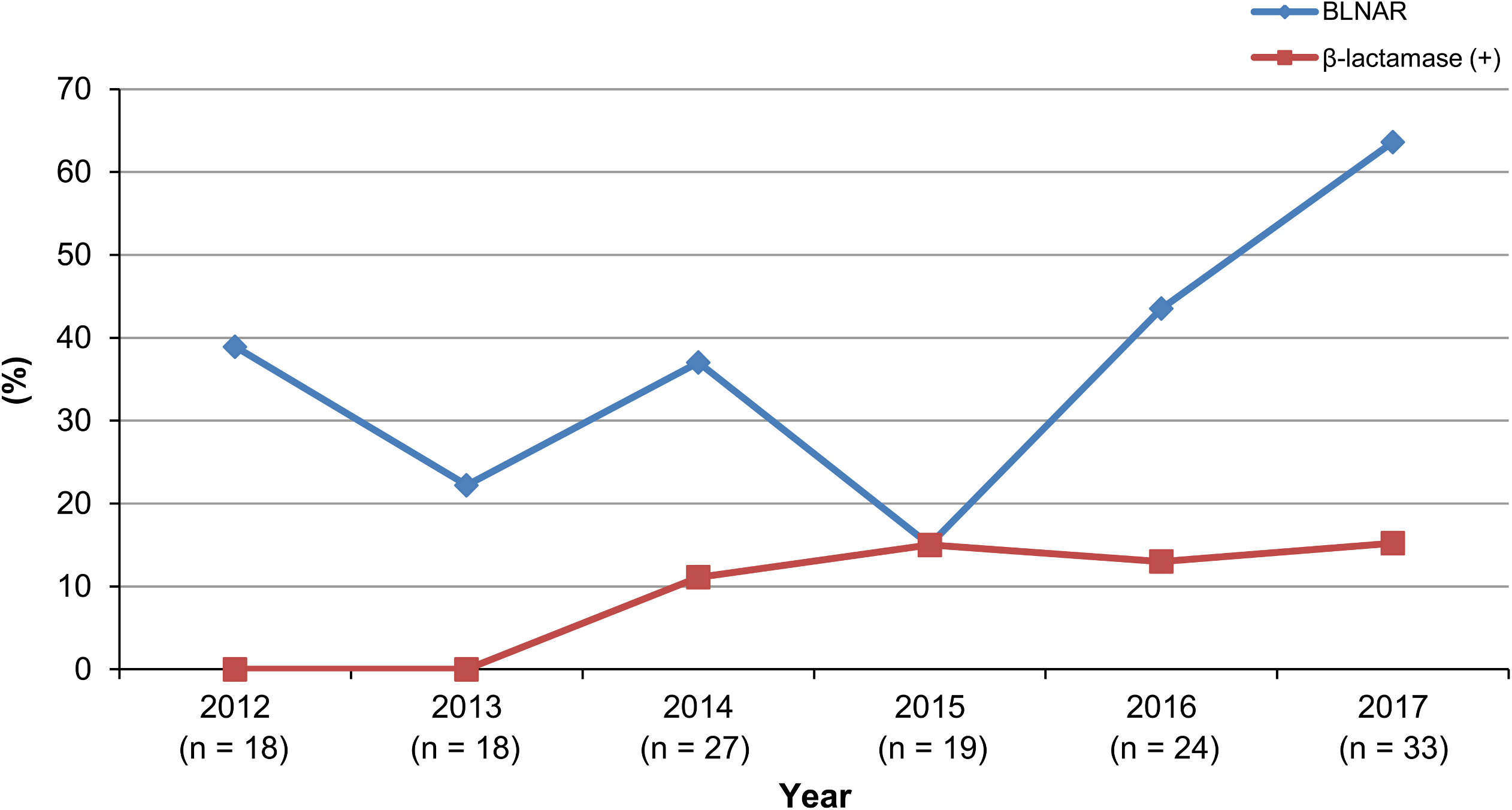
Percentages of BLNAR and β-lactamase-positive *Haemophilus influenzae* isolates in different years: Percentages of BLNAR isolates increased to be 15.0% in 2015, 43.5% in 2016, and 63.6% in 2017. β-Lactamase-producing isolates accounted for 15.2% in 2017.

### Susceptibility to quinolones

MIC values of various quinolone antibacterial agents measured with 144 *H. influenzae* isolates are shown for each year (Figure 2a–f). For all quinolones, not resistant strains as defined with MIC value of ≥2 µg/ml were found. Five quinolone antibacterial agents, except MXF, remained effective with MIC values of ≤0.03 µg/ml against approximately 90% of isolates each year (Figure 2b–f). For STFX in particular, no strains showing MIC values of ≥0.12 µg/ml were detected during the study period (Figure 2f).

**Figure 2a-f.**
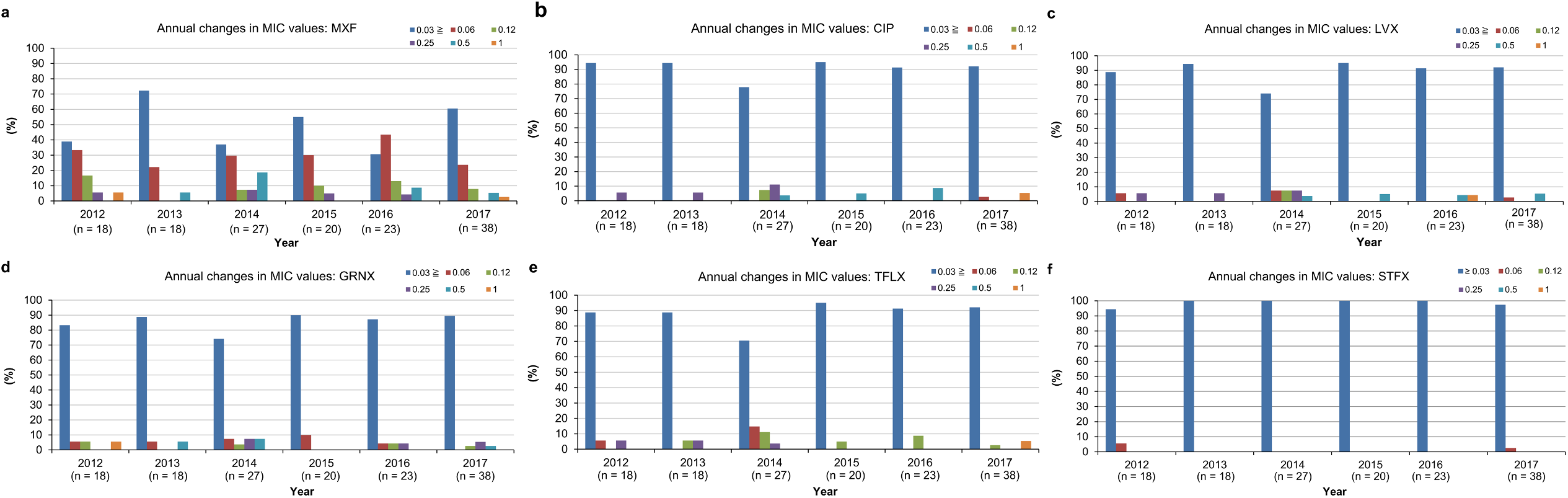
MIC values of fluoroquinolones for *Haemophilus influenzae* isolates in different years a: MXF MICs in different years; Isolates showing MIC value of ≥0.5 were detected, those with MIC value of ≤0.03 accounted for ≤80%, and the susceptibility decreased every year since 2013, except for 2015. b: CIP MICs in different years: The susceptibility is mostly maintained, although isolates with MIC value of ≥0.5 were detected since 2014. c: LVX MICs in different years: The susceptibility is mostly maintained, although isolates with MIC value of ≥0.5 were detected since 2014. d: GRNX MICs in different years: The susceptibility is mostly maintained, although isolates with MIC value of 0.5 were detected in 2013, 2014, and 2017. e: TFLX MICs in different years: The susceptibility is mostly maintained, although isolates with MIC value of 1 were detected in 2017. f: STFX MICs in different years; The susceptibility is nearly absolutely maintained, as all isolates had MIC value of ≤0.06.

### Comparison of susceptibility to MXF and other quinolones

Meanwhile, for MXF, a certain number of isolates with MIC values of ≥0.5µg/ml were detected every year since 2013 except 2015, and isolates with MIC value of ≤0.03 µg/ml accounted for less than 80% of isolates every year (Figure 2a). The MXF MIC value and relative MIC values of other quinolones against the same strain were compared. In an MIC range of 0.25–1.0 µg/ml, 64.7%, 70.6%, 76.5%, 88.2%, and 100% of the 144 strains tested showed MIC values of CIP (Figure 3a), LVX (Figure 3b), garenoxacin (GRNX), tosufloxacin (TFLX), and STFX (not shown in Figure) which were lower than that of MXF. This series of results demonstrated that the antibacterial potency of MFLX was clearly lower than that of any other antibacterial agent tested.

**Figure 3 a-b.**
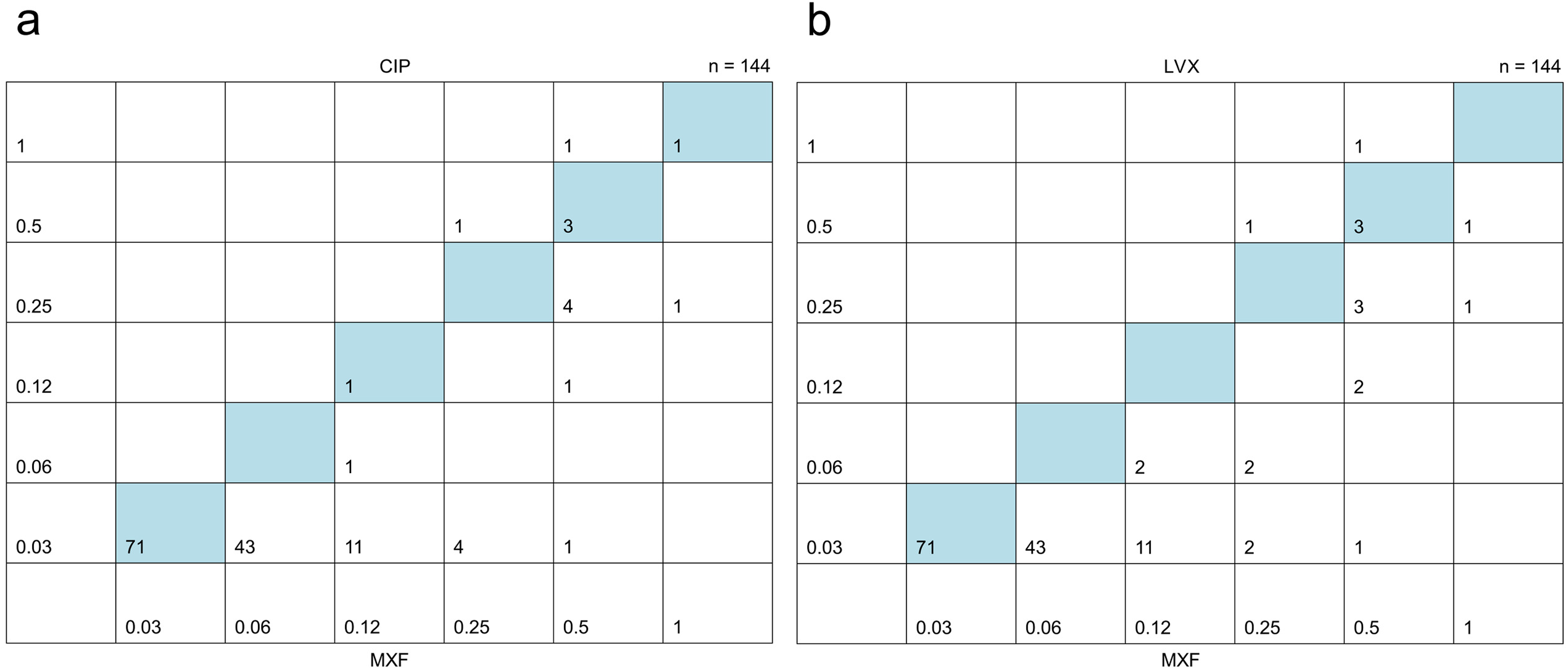
Correlation between MIC value of MXF and MIC values of other fluoroquinolones against *Haemophilus influenzae* (comparison in the same isolates) a: Correlation between MIC values of CIP and MXF: In a range of 0.25 ≤ MIC ≤ 1.0, MIC value of MXF was higher than that of CIP in 64.7% of isolates. b: Correlation between MIC values of LVX and MXF: In a range of 0.25 ≤ MIC ≤ 1.0, MIC value of MXF was higher than that of LVX in 70.6% of isolates.

### Amino acid mutations in QRDRs of GyrA and ParC

We investigated amino acid mutations in QRDRs of GyrA and ParC in 17 strains with low susceptibility to MXF (MIC ≥ 0.25 µg/ml) like a report described by Shoji et al (13). Genomic DNA of *H. influenzae* was purified, and any gene encoding QRDRs was PCR-amplified. Based on the nucleotide sequences thus obtained, amino acid sequences were analyzed, and amino acid mutations in QRDRs were identified. Mutations were found in 11 strains, and detected every year except 2015 (Table 1). Detection rates were 11.1% in 2012, 5.6% in 2013, 14.8% in 2014, 8.3% in 2016, and 5.3% in 2017 (total 7.6%). Only one mutation was found in GyrA (Ser84Leu) in 8 strains or in ParC (Ser84Ile or Gly82Arg) in 2 strains, and one mutation each in GyrA (Ser84Leu) and ParC (Ser84Ile). Both strains with the Ser84Ile mutation had MXF MIC ≥ 0.5 µg/ml, CIP MIC = 1 µg/ml, GRNX MIC ≥ 0.25 µg/ml, LVX MIC = 0.5 µg/ml, and TFLX MIC = 1 µg/ml, which indicated reduced susceptibility.

**Table 1.** Association between low quinolone susceptibility and QRDR mutations

| Strain |  |  |  |  |  |  | Mutation |  |
| --- | --- | --- | --- | --- | --- | --- | --- | --- |
|  | MFLX | CPFX | GRNX | LVFX | TFLX | STFX | GyrA | ParC |
| 2012—1 | 1 | 0.25 | 1 | 0.25 | 0.25 | 0.06 | Ser84Leu | — |
| 2012—2 | 0.25 | 0.03 | 0.12 | 0.06 | 0.03 | 0.03 | — | Gly82Arg |
| 2013—1 | 0.5 | 0.25 | 0.5 | 0.25 | 0.25 | 0.03 | Ser84Leu | — |
| 2014—1 | 0.5 | 0.25 | 0.5 | 0.25 | 0.25 | 0.03 | Ser84Leu | — |
| 2014—2 | 0.5 | 0.25 | 0.5 | 0.12 | 0.12 | 0.03 | Ser84Leu | — |
| 2014—3 | 0.5 | 0.25 | 0.25 | 0.25 | 0.12 | 0.03 | Ser84Leu | — |
| 2014—4 | 0.25 | 0.12 | 0.25 | 0.12 | 0.06 | 0.03 | Ser84Leu | — |
| 2014—5 | 0.5 | 0.5 | 0.12 | 0.5 | 0.12 | 0.03 | — | — |
| 2014—6 | 0.25 | 0.03 | 0.06 | 0.06 | 0.06 | 0.03 | — | — |
| 2014—7 | 0.25 | 0.03 | 0.06 | 0.03 | 0.06 | 0.03 | — | — |
| 2015—1 | 0.25 | 0.5 | 0.06 | 0.5 | 0.12 | 0.03 | — | — |
| 2016—1 | 0.5 | 0.5 | 0.25 | 1 | 0.12 | 0.03 | Ser84Leu | — |
| 2016—2 | 0.5 | 0.5 | 0.12 | 0.5 | 0.12 | 0.03 | Ser84Leu | — |
| 2016—3 | 0.25 | 0.03 | 0.06 | 0.03 | 0.03 | 0.03 | — | — |
| 2017—1 | 1 | 1 | 0.5 | 0.5 | 1 | 0.06 | — | Ser84Ile |
| 2017—2 | 0.5 | 1 | 0.25 | 0.5 | 1 | 0.03 | Ser84Leu | Ser84Ile |
| 2017—3 | 0.5 | 0.03 | 0.12 | 0.03 | 0.03 | 0.03 | — | — |

BLNAR and amino acid mutations in QRDRs. Of the 11 isolates with any amino acid mutation, 5 (2012-2, 2014-1, 2014-3, 2017-1, and 2017-2) were BLNAR, and all but 2012-2 exhibited reduced susceptibility to fluoroquinolones except for STFX (Table 1).

## Discussion

In Japan, no Hib strains have been detected in children with invasive infections after Hib vaccines were introduced in 2008 (14). However, the frequency of isolation of BLNAR strains has not decreased; It is rather high in some areas, with 33.5% reported in the respiratory area in 2010 (15), 35.8% in the otorhinolaryngology area in 2012 (3), and 46.7% in the pediatric respiratory area in 2012 (16). Additionally, the present study showed that the frequency of isolation of BLNAR strains increased to 63.6% in 2017 (Figure 1).

Fluoroquinolone resistance rates of *H. influenzae* previously studied in Japan were extremely low, and the 2010 respiratory specimen surveillance report also stated that the resistance to CIP and LVX was found in 0.5% and STFX resistance was found in 0% of isolates (15). STFX is characterized by the strong inhibitory potencies against both DNA gyrase and topoisomerase IV (17). The well-preserved STFX sensitivity owing to this characteristic was consistent with that reported in other reports (8–13) and the present findings.

Meanwhile, the MXF resistance rate in *H. influenzae* isolates has been reported to be 1.1% in 2010 (15); however, it is noteworthy that low susceptibility to MXF was observed for 11.8% (17 of 144) of isolates collected from 2012 to 2017 in the present study. This is likely to be attributable to the fact that MXF can be orally administered in Japan, whereas injectable formulations are available in Europe and the United States. The mutant prevention concentration of MXF for *H. influenzae* has been reported to be 0.125–0.25 µg/ml (18). However, the resistance may be acquired if the MXF level in blood does not sufficiently increase because an antacid or laxative, for example, is concomitantly used with oral MXF. Furthermore, the drug resistance epidemic status in the area should be understood.

Additionally, MIC values of LVX and GRNX, which are frequently used in Japan, and MIC value of CIP, which is often used against *Pseudomonas aeruginosa*, also tended to be elevated. In 2017, two isolates with TFLX MIC = 1 µg/ml (one from a child) were identified (Figure 2e). After the recent approval for treatment of respiratory infections in children as an additional indication in Japan, opportunities of prescribing TFLX are increasing, and there is a concern that this might lead to an increase in the prevalence of TFLX resistance.

Considering that MIC value of MXF is relatively higher than the MIC values of other quinolones (Figure 3a,b) and two of six isolates with MXF MIC value of 0.25 µg/ml had amino acid mutations in QRDRs (Table 1), MIC values of MXF appear to be more useful than those of other quinolones as a reference to find bacterial isolates with reduced susceptibility to quinolone antibacterial agents. Therefore, we believe that MXF is suited for MIC assays to screen bacterial isolates with reduced susceptibility to fluoroquinolones, particularly in Japan. Furthermore, considering the reports by Shoji (13) and Perez (6), we propose that an MXF MIC value of 0.25 µg/ml is an appropriate threshold to screen bacterial isolates for amino acid mutations in QRDRs. QRDR mutations are found more frequently at Ser84 and Asp88 of GyrA gene and at Ser84, Glu88, and Gly82 of ParC gene than at other positions (5,8,13). In the case of *H. influenzae*, previous studies have demonstrated that MICs increase when strains have mutations at Ser84 in GyrA QRDRs or at Gly82 or Ser84 in ParC QRDRs (5,7,13). In this study, we analyzed 17 isolates showing MXF MIC value of ≥0.25 µg/ml and found that 9 isolates had a leucine mutation at Ser84 of GyrA gene (Table 1), a finding that is consistent with previous reports. Considering the reports on *Escherichia coli*, *Klebsiella pneumoniae* (19), *Enterococcus faecalis* (20), and other bacteria, this finding suggests that GyrA is a likely primary target of quinolones for their antimicrobial activity similary, against *H. influenzae*.

In this study, 2012-2 and 2017-1 isolates were found to have mutations only in ParC (Table 1). Related to this result, Shoji et al obtained a mutant with the mutation only at Ser84 of ParC gene among hot spots in a resistance induction experiment (13). The MXF MIC for this mutant was 0.5 µg/ml, indicating low susceptibility. This induction experiment showed that Ser84 is an important mutation site for *H. influenzae* to acquire quinolone resistance, supporting the finding in the clinical isolates in the present study.

While MIC values have been reported to increase as the number of QRDR mutations increases (13–21), only one isolate was found to have two mutations in the present study. The isolate 2017-2 showed reduced susceptibility to all fluoroquinolones except for STFX, a finding supporting the above notion (Table 1).

Isolate 2014-5 showed a high MIC value (0.5 µg/ml) for not only MXF but also CIP and LVX, although this isolate had no mutations in QRDRs of both GyrA and ParC enzymes (Table 1). A factor presumably involved in this is a drug efflux pump (acrAB) (21–22) or *H. influenzae*-specific porin (23–24), the protein structure of which is similar to that of porin (OmpF); however, solid evidence was not obtained from the present study, and this point remains to be addressed in future studies.

It is difficult to predict whether the frequency of quinolone-resistant *H. influenzae* isolates will increase in the future. Nevertheless, we consider that an outbreak of quinolone-resistant *H. influenzae* is likely to occur because the population aged >65 years is a risk factor for this resistance (7), quinolone-resistant *E. coli* is increasing rapidly over recent years, and quinolone-resistant *E. coli* isolates also have mutations in DNA gyrase and topoisomerase IV (19). In 2015, a global action plan on antimicrobial resistance (AMR) was adopted by the World Health Assembly, and member countries developed national action plans. In Japan, a milestone is to halve the amount of use of broad-spectrum oral antibacterial agents, including oral fluoroquinolones by 2020. Therefore, monitoring the trend of occurrence of fluoroquinolone resistant *H. influenzae* isolates is likely to have a substantial meaning to AMR countermeasures.

In clinical practice, the presence of a drug resistance gene does not necessarily indicate that the isolate has a reduced susceptibility to that antibacterial agent (i.e., silence mutations) (25). However, acquisition of quinolone resistance is facilitated by the use of quinolones. Therefore, the emergence of quinolone-resistant bacteria can be prevented and future medical care against infectious diseases would benefit if we can eliminate the unnecessary use of antibacterial drugs by promoting proper use of antibacterial drugs. Limitations of this study include that we could not follow clinical courses of cases in which isolates with low quinolone susceptibility were found and that we could not determine the resistance status in other institutions. Close monitoring of changes in drug resistance and clinical characteristics should be jointly performed at multiple sites.

## Materials and Methods

### Bacterial strains

A total of 144 *H. influenzae* strains that had been clinically isolated at the Showa University Hospital between 2012 and 2017 and stored in our laboratory were used in this study.

### β-Lactamase-producing ability test

Each bacterial strain was cultured on chocolate agar medium (Nippon Beckton-Dickinson, Tokyo, Japan) at 35°C in 5% CO_2_ for 22 hours (8), and then nitrocefin (Kantokagaku, Tokyo, Japan) was added in drops to *H. influenzae*; the strain was considered to be capable of producing β-lactamase when the red color developed (12).

### Drug susceptibility assay

Susceptibility to antibacterial agents was measured by the Clinical and Laboratory Standards Institute broth microdilution method using 96-well plates (11). As a basal medium, the liquid medium added to each well was HTM medium (Eiken, Tokyo, Japan) and was supplemented with Hematin (15 µg/ml) and NAD (15 µg/ml) (Eiken Chemical, Class I bacteria test series, frozen plates). Each strain was cultured on a chocolate agar medium at 35°C in 5% CO_2_ for 22 hours, and then a bacterial suspension was prepared at a concentration adjusted to be similar to that of McFarland standard (bioMèrieux) 0.5 using physiological saline. Finally, each well was inoculated with a bacterial suspension (5 × 10^5^ CFU/ml). The plate was cultured at 35°C in 5% CO_2_ for 22 hours, and MIC values of test drugs for these bacteria were measured. The following test drugs were used:

> aminobenzylpenicillin (ABPC), sulbactam/ampicillin (SAM), clavulanic acid/amoxicillin (AMC), levofloxacin (LVX), ciprofloxacin (CIP), garenoxacin (GRNX), moxifloxacin (MXF), tosufloxacin (TFLX), and sitafloxacin (STFX).

### Analysis of QRDR amino acid sequences

*H. influenzae* strains showing MXF MIC value of ≥0.25 µg/ml in the drug susceptibility assay were selected as test strains and allowed to grow. From bacterial cells of the test strains, genomic DNA was extracted using Sepa-Gene (EIDIA Co., Ltd., Tokyo, Japan), allowed to aggregate by adding 99.5% alcohol, centrifuged at 8000 ×*g* to precipitate, and then washed with 70% alcohol. Using the resulting DNA sample as a template, genes encoding QRDRs were amplified by PCR. For amplification, a mixture solution of *Taq* polymerase (Roche, Mannheim, Germany) and primer (26) was used in a T100 thermocycler (BIORAD, CA, USA) under PCR conditions similar to those described by Shoji et al (13).

Next, the PCR product was subjected to purification/separation and extraction as follows. The entire PCR product was subjected to electrophoresis in 2% agarose gel (TAKARA, Shiga, Japan) under 120 V/h for 1 hour in Tris-borate-EDTA (NIPPON GENE, Tokyo, Japan) buffer. Then, the gel was stained with ethidium bromide to detect the target PCR product bands (GyrA: 375 bp, ParC: 370 bp), and the detected bands were excised. Thereafter, from the excised gel segments, DNA was extracted and purified using a GenElute Minus EtBr Spin Column (SIGMA-ALDRICH, St. Louis, USA) and concentrated with ethanol. For direct sequencing using the resulting DNA samples as templates, PCR amplification was performed using Big Dye Terminator v1.1 cycle sequencing kit (Applied Biosystems Japan, Tokyo, Japan). The PCR primers described above were used here as sequence primers.

The resulting sequencing solutions were analyzed using an ABI PRISM 310 genetic analyzer (Applied Biosystems, Foster City, CA), and nucleotide sequences of the target gene regions were determined. Based on the nucleotide sequence data obtained, amino acid sequences were analyzed using GENETYX Ver. 8 (GENETYX, Tokyo, Japan) to find amino acid mutations in QRDRs.

## Acknowledgments

This work was supported by Kazuhisa Ugajin. We thank Zhao WH for providing the nitrocefin reagent.

## Transparency declarations

We have no Conflict of Interest to declare. This research received no specific grant from any funding agency in the public, commercial, or not-for-profit sectors.

